# CRISPR-Cas9 editing induces Loss of Heterozygosity in the pathogenic yeast *Candida parapsilosis*

**DOI:** 10.1101/2022.08.15.504067

**Authors:** Lisa Lombardi, Sean A. Bergin, Adam Ryan, Geraldine Butler

**Author notes:** Corresponding author Lisa Lombardi, School of Biomolecular and Biomedical Science, Conway Institute, University College Dublin, Belfield, Dublin 4.

## Abstract

Genetic manipulation is often used to study gene function. However, unplanned genome changes (including Single Nucleotide Polymorphisms (SNPs), aneuploidy and Loss of Heterozygosity (LOH)) can affect the phenotypic traits of the engineered strains. Here, we show that CRISPR-Cas9 editing can induce LOH in the diploid human pathogenic yeast *Candida parapsilosis*. We sequenced the genomes of ten isolates that were edited with CRISPR-Cas9 and found that the designed changes were present in nine. However, we also observed LOH in all isolates, and aneuploidy in two isolates. LOH occurred most commonly downstream of the Cas9 cut site and extended to the telomere in three isolates. In two isolates we observed LOH on chromosomes that were not targeted by CRISPR-Cas9. Two different isolates exhibited cysteine and methionine auxotrophy caused by LOH at a heterozygous site in *MET10*, approximately 11 and 157 kb downstream from the Cas9 target site, respectively. *C. parapsilosis* isolates have relatively low levels of heterozygosity. However, our results show that mutation complementation to confirm observed phenotypes is important even when using CRISPR-Cas9, which is now the gold standard of genetic engineering.

**IMPORTANCE:** CRISPR-Cas9 has greatly streamlined gene editing, and is now the gold standard and first choice for genetic engineering. However, we show that in diploid species extra care should be taken in confirming the cause of any phenotypic changes observed. We show that the Cas9-induced double strand break is often associated with loss of heterozygosity in the asexual diploid human fungal pathogen *Candida parapsilosis*. This can result in deleterious heterozygous variants (e.g. stop gain in one allele) becoming homozygous resulting in unplanned phenotypic changes. Our results stress the importance of mutation complementation even when using CRISPR-Cas9.

## INTRODUCTION

The opportunistic fungal pathogen *Candida parapsilosis* is a member of the CUG-Ser1 clade (1), and the second or third most frequently isolated *Candida* species (2). Unlike *Candida albicans, C. parapsilosis* has been associated with nosocomial infection outbreaks worldwide (3-6). Severe *C. parapsilosis* infections are a concern in intensive care units and neonatal intensive care units, where they are associated with neonatal mortality (7, 8).

Genetic manipulation of the diploid genome of *C. parapsilosis* is fundamental for studying the pathobiology of this species, which often cannot be directly extrapolated from *C. albicans*. Site directed mutagenesis in *C. parapsilosis* has been accomplished by homologous recombination using the *SAT1* flipper cassette containing a dominant selection marker (reviewed in (9)), or nutritional selection markers to replace the gene of interest in an auxotrophic background (10). More recently, CRISPR-Cas9 technology was adapted for the *C. parapsilosis sensu lato* complex (*C. parapsilosis, Candida orthopsilosis*, and *Candida metapsilosis*), which allows markerless editing in prototrophic strains to introduce homozygous or heterozygous mutations, to delete genes, and to tag proteins (11-18). We are currently using the plasmid-based CRISPR-Cas9 system, pCP-tRNA (12), to systematically disrupt genes in *C. parapsilosis* CLIB214.

Despite being a common approach for studying gene function, the construction of mutant strains can be associated with unplanned genome rearrangements, including aneuploidy and Loss of Heterozygosity (LOH) (19-25). Aneuploidy entails gain or loss of a chromosome or a chromosome segment, resulting in a change in gene dosage. LOH results in loss of genetic information from one of the two chromosome homologs. In *Candida albicans*, rates of both these events increase in response to stress (26, 27). Aneuploidy and LOH are also frequently found in strains of *C. albicans* that underwent manipulation in the laboratory (19-25). In fact, two commonly used transformation methods (lithium acetate and electroporation) promote changes in chromosome copy number, possibly by increasing chromosome non-disjunction, and exposure to heat preferentially triggers aneuploidy or LOH, depending on the length and intensity of the exposure (22). It is now known that deleting *ura3* in the *C. albicans* reference strain SC5314 to generate CAI-4 resulted in trisomy of Chr2 and/or Chr3 (19, 20), and additional manipulations generating double (*ura3/his1*) or triple (*ura3/his1/arg4*) auxotrophic strains resulted in further changes (20). Large tracts of LOH (up to 1330 kb) were also found on multiple chromosomes in other SC5314-derived strains (25).

Both aneuploidy and LOH can affect the phenotypic traits of a strain, which may be erroneously linked to the genome change that was intentionally introduced (28, 29). For example, in the process of deleting <u>S</u>erine <u>A</u>spartic <u>P</u>rotease genes in *C. albicans*, a *sap4/sap5/sap6* triple mutant was constructed in which LOH coincidentally resulted in the loss of the *SAP2-2* allele, present as a heterozygous locus (*SAP2-1/ SAP2-2*) on chromosome R. The inability of the resulting strain to use proteins as a sole nitrogen source was initially assumed to be due to the absence of Saps4-6, but was actually caused by the loss of the *SAP2-2* allele (23). Similarly, spontaneous LOH of the right arm of chromosome 3 determined sensitivity to the DNA-damaging agent methyl methane sulfonate (MMS) depending on which *MBP1* allele was retained (24).

Although CRISPR-Cas9 revolutionized the landscape of gene editing, an increasing number of studies shows that this technology can also induce aneuploidy and LOH in eukaryotic cells (30-33). A recent study found that CRISPR-Cas9-generated *C. albicans* strains contain numerous unwanted genomic changes, albeit to a lesser extent than the strains generated with other methods (31). To evaluate the extent of LOH and aneuploidy induced by gene editing in *C. parapsilosis*, we sequenced the genomes of ten CRISPR-Cas9 genetically modified strains (nine edited strains and one gene deletion). We found that LOH occurs frequently, and it is more common on chromosomes targeted by Cas9. We also show that LOH on chromosome 8 can result in cysteine/methionine auxotrophy. Overall our study stresses the importance of confirming a direct relationship between genotype and phenotype when using CRISPR-Cas9, especially when working with diploid species.

## RESULTS

### Gene manipulation is associated with unplanned genome changes in *C. parapsilosis*

We previously described the construction of homozygous deletions of 73 transcription factors and 16 protein kinases in *C. parapsilosis*, generated by using homologous recombination to replace both alleles at a single locus with either *CdHIS1* or *CmLEU2* (10). We are currently using a plasmid-based CRISPR-Cas9 system (12) to disrupt additional genes by introducing 11 bases containing stop codons in all open reading frames and a unique barcode (Fig. 1).

**Fig 1.**
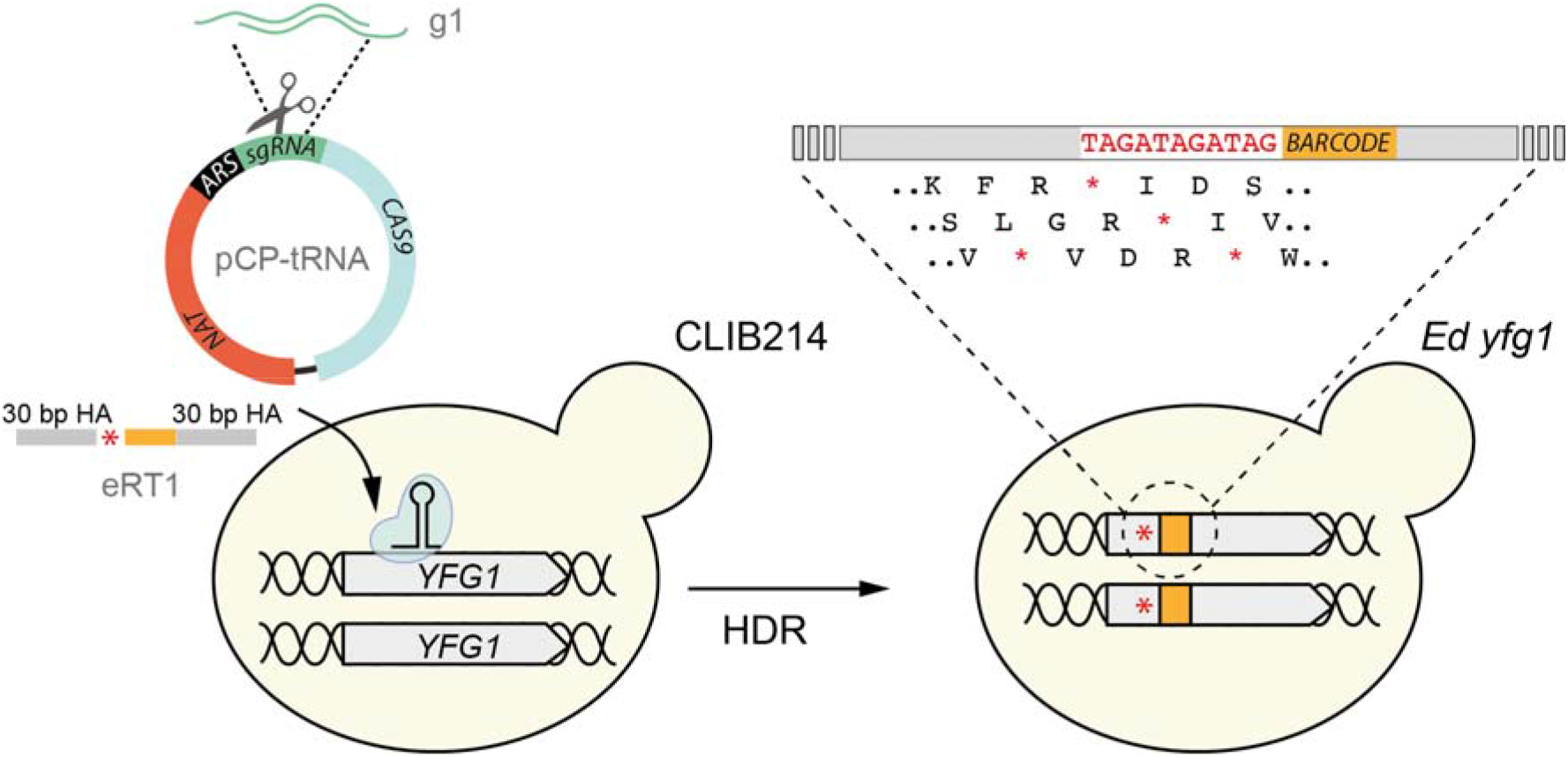
Workflow for the construction of CRISPR-Cas9 edited strains in *C. parapsilosis* CLIB214. Guides were designed to target Cas9 cleavage to the first third of each open reading frame (*YFG*, Your Favourite Gene). The Repair Templates (e.g. eRT1) contain 11 bp with stop codons in each of the 3 possible reading frames (red asterisk) and a unique barcode (in orange), flanked by two short Homology Arms (HAs) to drive Homology Directed Repair (HDR). The short dsDNA guides (e.g. g1) were cloned into the pCP-tRNA plasmid, which harbours *CAS9*, the Autonomously Replicating Sequence (*ARS*), and a nourseothricin marker cassette (*NAT*) (12). Simultaneous transformation of the pCP-tRNA (e.g. pCP-tRNA1) targeting a gene (e.g. *YFG1*) and the corresponding RT (e.g. eRT1) results in the insertion of premature stop codons (red asterisk) in both alleles by HDR (e.g. Ed yfg1).

To determine the level of unplanned changes that occur during CRISPR-Cas9 editing, we sequenced the genomes of nine strains that were edited in nine different genes. The names of the edited strains (e.g. e301940) reflect the target gene (e.g. *CPAR2_301940*). *C. parapsilosis* is on average less heterozygous than *C. albicans* (0.1-0.4 SNP/kb versus 3 SNP/kb, respectively) (34, 35). To maximize the chances of detecting LOH, we selected strains in which the target gene was followed by regions containing at least some heterozygous sites. Many of these are on Chromosome 8; as a consequence, strains edited in genes located on chromosome 8 were overrepresented in the sequenced strains (Table 1, Fig. 2).

**Table 1.** LOH in CLIB214-derived strains modified using CRISPR-Cas9 technology. Grey colour shows LOH at the chromosomes targeted by CRISPR-Cas9.

| Strain | Site of edit (Cas9 cut site) | LOH Coordinates<sup>1</sup> | LOH range |
| --- | --- | --- | --- |
| e101740 | Chr1: 386,438 | Chr5: 104,538 - 661,431 | 557 kb |
| e205070 | Chr2: 1,028,668 | Chr2: 1,022,177 - 1,024,638 | 2.5 kb |
| e301780 | Chr4: 417,148 | Chr4: 417,081 | Single variant |
| e301940 | Chr3: 898,895 | Chr2: 1,540,134 - 1,619,139 | 79 kb |
| e802210 | Chr8: 504,595 | Chr8: 142,590 - 567,069 | 424 kb<sup>2</sup> |
| e804640 | Chr8: 1,033,704 | Chr8: 142,590 – 1,041,458 | 899 kb<sup>2</sup> |
| e804830 | Chr8: 1,080,807 | Chr8: 142,590 – 1,078,006 | 935 kb<sup>2</sup> |
| e804940 | Chr8: 1,097,623 | Chr8: 1,091,412 | Single variant |
| e805700 | Chr8: 1,290,224 | Chr8: 1,292,562 - 1,453,338 | 161 kb |

<sup>1</sup> The positions of the first and last variants in the LOH block.
<sup>2</sup> Likely reaches the telomere.

**Fig. 2.**
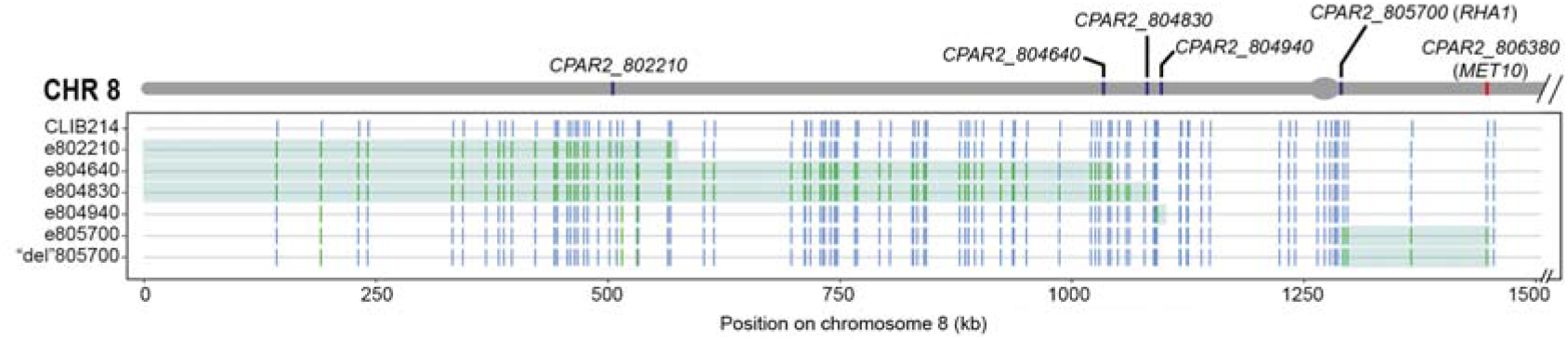
CRISPR-Cas9 induces LOH on chromosome 8. The grey bar on top represents chromosome 8, with target genes in blue and *MET10* in red. The centromere is indicated by the grey oval. For each strain, heterozygous sites are shown in blue and homozygous sites in green, with respect to the reference genome (CDC 317). LOH tracts are shaded in green. The final 593 kb of the chromosome is not shown. CLIB214 = *C. parapsilosis* CLIB214; e802210 = stop codons inserted in *CPAR2_802210* (and so on for the other strains and genes); “del”805700 indicates a strain in which Cas9 likely cut at *CPAR2_805700* but the repair template was not incorporated.

The designed edit was present in all strains. Two edited strains (e205070 and e804640) contained three copies of chromosomes 6 and 5, respectively (Fig. S1). We observed LOH in all the strains (Table 1). In two isolates only one single heterozygous variant was lost (e301780 and e804940), located 67 bp and 6 kb from the target cut site, respectively. In the remaining seven strains, LOH ranged from 2.5 to more than 900 kb, and most likely to the telomere in three isolates (Table 1, Fig. 2).

### CRISPR-Cas9 induces Loss of Heterozygosity in *C. parapsilosis*

LOH occurred more frequently on the chromosome on which the target gene was located (6/9 strains), suggesting that it may be induced by the Cas9-induced double strand break (DSB). This is shown in detail for chromosome 8 (Fig. 2); LOH starts at the Cas9 cut site, or near the cut site strongly suggesting that LOH results from homology-based repair mechanisms (e.g. Break-induced Replication, BIR (36, 37) triggered by the DSB. In three strains we also observed LOH on chromosomes that were not targeted by CRISPR-Cas9 (Table 1).

### LOH on chromosome 8 can result in cysteine/methionine auxotrophy

While testing the phenotypes of edited strains, we noticed that one isolate containing stop codons in *CPAR2_805700* (*RHA1*) on chromosome 8, failed to grow in the absence of the sulfur containing amino acids cysteine or methionine (Fig.3). This strain (e805700) was generated by targeting *RHAI* with the plasmid pCP-805700. *RHA1* encodes a zinc cluster transcription factor that acts as a positive filamentation regulator (38) and has never been shown to be involved in sulfur metabolism.

To further characterize the observed phenotype, we attempted to use CRISPR-Cas9 to delete *RHA1*, using plasmid pCP-805700. The deletion strategy was unsuccessful: the resulting strains (called “del”805700) contained a functional *RHA1* gene, but they again failed to grow in the absence of cysteine or methionine (Fig. 3). Comparing the genomes of *C. parapsilosis* CLIB214, e805700 (edited *RHA1*), and “del”805700 showed that both CRISPR-Cas9 manipulated strains had a stretch of LOH on chromosome 8 (1,292,562 – 1,453,338) extending from *RHA1* to *CPAR2_806380* (*MET10*) (Fig 3). In *C. parapsilosis* CLIB214 one allele of *MET10* contains a premature stop codon (Table S1). In both e805700 and “del”805700, LOH resulted in homozygosis at this site that caused loss of function of *MET10*; both alleles contain a C2600A SNP converting Ser867 into a stop codon. *MET10* encodes one of the catalytic subunits of the sulfite reductase, which reduces sulfite to sulfide in the sulfur assimilation pathway (Fig. 3B). This enzyme is a heterotetramer composed of 2 α and 2 β subunits (α2β2), encoded by *MET10* and *MET5*, respectively. The lack of a functional *MET10* results in a non-functional sulfite reductase, and this shuts down the pathway before the production of homocysteine, *de facto* preventing the cells from producing cysteine and methionine (Fig. 3B) (39).

**Fig 3.**
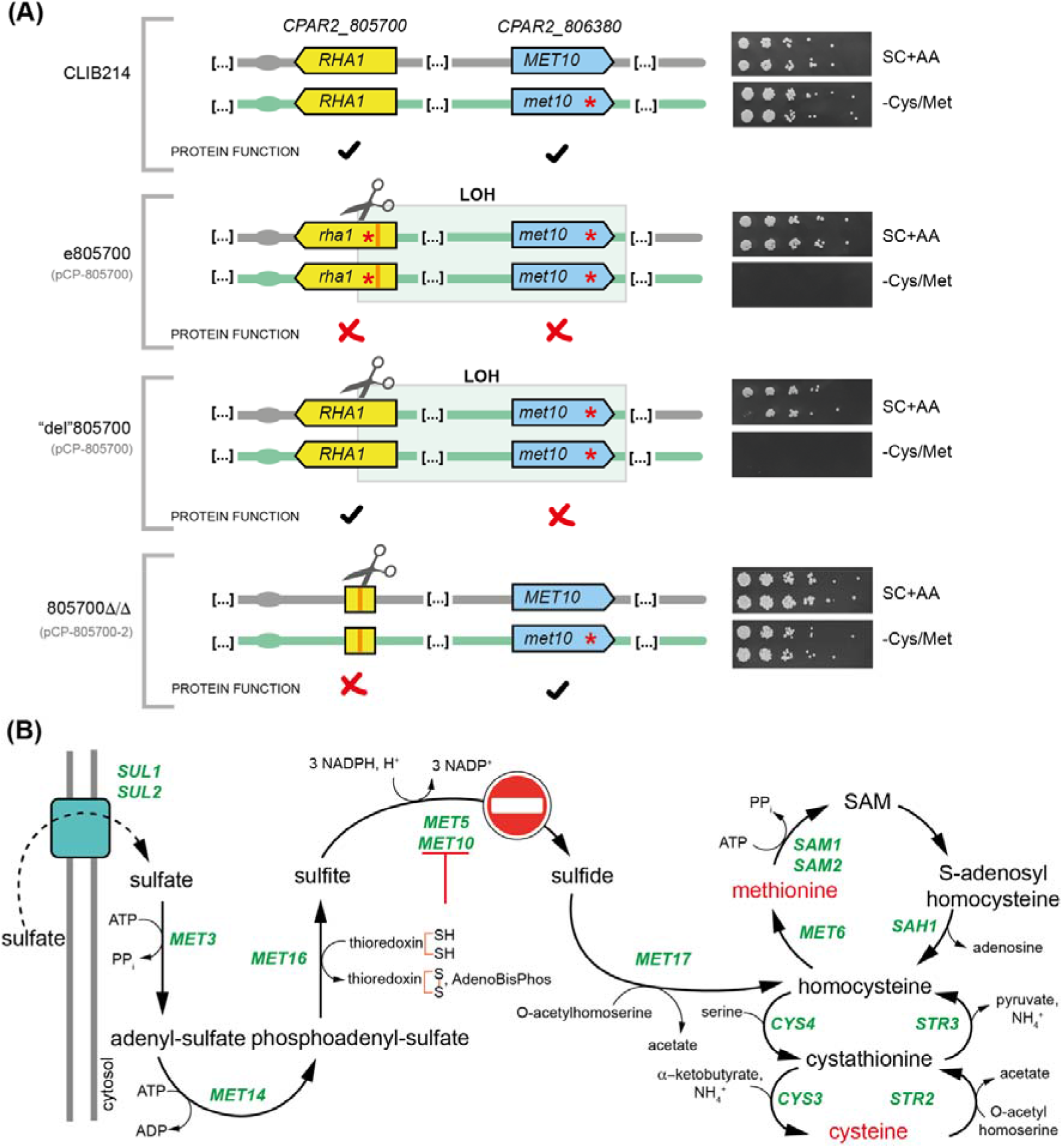
LOH-induced loss of function of the sulfite reductase Met10 causes cys/met auxotrophy. (A) The diagrams on the left show the presence of functional alleles of *RHA1* and *MET10* or of alleles containing stop codons (red asterisks). The plasmids used to target *CPAR2_805700* (*RHA1*) are indicated in grey underneath the name of each strain. Note that 805700Δ/Δ was generated using a different guide than e805700 and “del”805700. Images on the right show the growth of serial dilutions of two biological replicates of each strain on SC with all amino acids, or SC missing cysteine and methionine. In *C. parapsilosis* CLIB214, one allele of *CPAR2_806380* (*MET10*) has a premature stop codon. Strains that have undergone homozygosis at this position, generating two *MET10* alleles with stop codons, fail to grow in the absence of cysteine and methionine (e.g. e805700 and “del”805700). Strain 805700Δ/Δ contains a partial deletion of *RHA1* with no LOH at *MET10*. Cas9 target sites are indicated with a scissors icon. (B) Sulfur assimilation pathway in *Saccharomyces cerevisiae* (39). In the absence of cysteine and methionine, yeast cells can import sulfate from the extracellular environment, and then gradually reduce it to sulfide, which is then incorporated into homocysteine. Homocysteine is then funneled into the methyl cycle to produce methionine, and into the transsulfuration pathway to synthesize cysteine. *MET10* and *MET5* encode the two catalytic subunits (α and β, respectively) of the α2β2 heterotetrameric enzyme sulfite reductase. Loss of function of Met10 shuts down the pathway, thus making cells auxotrophic for cys/met. SAM: S-adenosyl methionine.

To confirm that *RHA1* does not play a role in sulfur metabolism, we used a different guide RNA to delete the gene (pCP-805700-2, Fig. 3). All 11 transformants obtained were deleted for *RHA1*, and they could grow in the absence of cysteine and methionine (Fig. S2). By PCR amplifying and sequencing the fragment of *MET10* containing the heterozygous variant, we showed that all the isolates retained both alleles (wild type and stop codon), and that they had a functional copy of Met10 like the parental strain CLIB214 (Fig. S2).

In an independent experiment, we noticed that disrupting *CPAR2_806320* (*KIS1*) on chromosome 8 also resulted in cys/met auxotrophy (e806320, Fig. 4). In *C. albicans*, Kis1 is one of the two β subunits of the Snf1p complex, and *KIS1*-deficient mutants fail to grow on many alternative carbon sources (40). As with *RHA1*, no role in sulfur metabolism had been described for *KIS1*. The *C. parapsilosis* CRISPR-Cas9 edited e806320 strain cannot grow on lactose, sodium acetate, ethanol, or glycerol as sole carbon sources, similar to *C. albicans*, but it also fails to grow in the absence of cysteine and methionine (Fig. 4). In contrast, a strain in which both *KIS1* alleles were deleted by homologous recombination (kis1Δ/Δ, (10)), fails to use alternative carbon sources, but is a cysteine and methionine prototroph (Fig. 4). The genome of e806320 was not sequenced. However, PCR amplification and Sanger sequencing showed that the edited strain e806320 has undergone LOH resulting in stop codons at both *MET10* alleles (Fig. 4). In the deleted strain kis1Δ/Δ one wild type allele of *MET10* is retained (Fig. 4).

**Fig 4.**
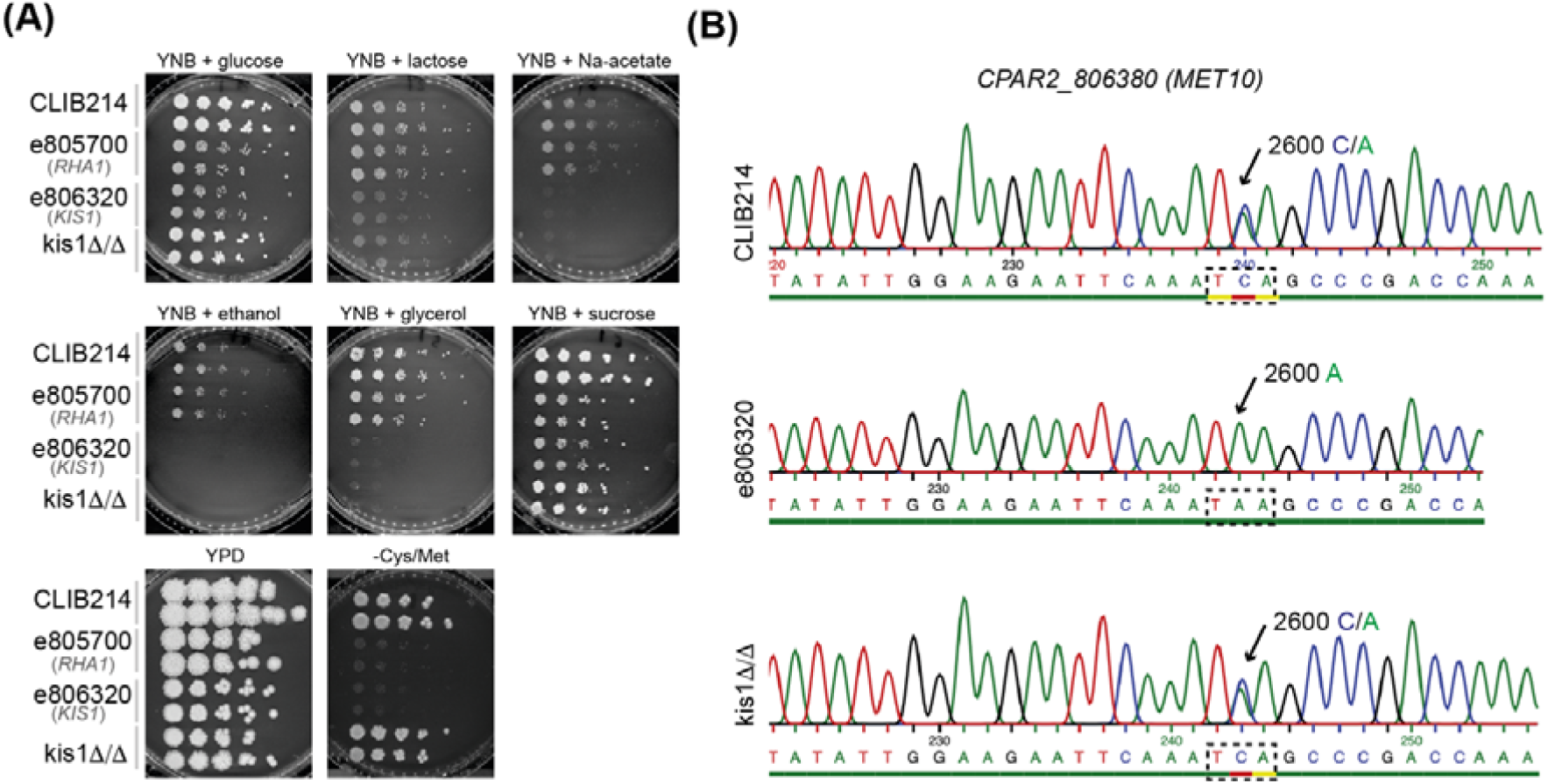
Kis1 is required for utilization of alternative carbon sources, but not sulfur metabolism. A.Growth of parental strain CLIB214 and its derivatives, e805700 (edited *RHA1*), e806320 (edited *KIS1*), and kisΔ/Δ (deleted *KIS1* by allele replacement, (10). The gene targeted by Cas9 is indicated in grey underneath the name of the edited strains. Editing or deleting *KIS1* reduces growth on lactose, sodium acetate, ethanol, and glycerol as sole carbon sources. However, only the edited strain is also auxotrophic for cys/met, similar to the strain edited in *RHA1* (e805700). (B) Sequencing of the *CPAR2_806380* (*MET10*) locus shows that there is a heterozygous site (C/A in position +2600) in both CLIB214 and kis1Δ/Δ. However e806320 is homozygous at this position (2600A) showing that both *MET10* alleles in this strain have stop codons.

### CRISPR-Cas9 associated LOH may result in other unexpected phenotypes

Our results show that CRISPR-Cas9-induced LOH could cause unplanned phenotypes in edited strains. To determine how widespread the phenomenon could be, we identified all heterozygous sites in *C. parapsilosis* CLIB214 where homozygosis is predicted to result in deleterious phenotypes. We found 71 heterozygous variants in 67 genes, including 13 stop-gains, 6 frameshift mutations, and 52 potentially deleterious non synonymous amino acid changes (SIFT score < 0.05, Table S1). 51 genes had orthologs in *Saccharomyces cerevisiae* (Table S1). The variants are not equally distributed throughout the eight chromosomes: 26, including 5 stop codons, are located on chromosome 8, reflecting the higher heterozygosity of this chromosome. Overall, the effect of a non-synonymous amino acid change is often not easy to predict, and therefore the 52 variants of this type may overestimate the number of potentially deleterious alleles. By contrast, a frameshift mutation or a stop-gain mutation are highly likely to disrupt protein function.

## DISCUSSION

Although CRISPR-Cas9 streamlined gene editing of asexual diploid species like *C. parapsilosis*, the genome-scale aftermath of CRISPR-Cas9 editing has not been studied in this fungal pathogen, and studies in any *Candida* species are rare (31). We found some unplanned changes on chromosomes that were not directly targeted by Cas9, including trisomy of chromosomes 5 or 6, and LOH (Table 1, Fig. S1), suggesting that transformation is a mutagenic process in *C. parapsilosis*, or indeed that CRISPR-Cas9 may induce LOH in places others than the targeted loci. Transformation-induced aneuploidy and LOH are common in laboratory strains of *C. albicans* (19-25). Using a transient CRISPR-Cas9 system to simultaneously integrate two nutritional markers (*ARG4/HIS1*) in the two alleles of targeted genes was also associated with LOH events that were not on the targeted chromosomes (31). These genomic changes sometimes result in phenotypic changes, such to nitrogen utilization (23), tolerance to DNA-damaging agents (24), and virulence *in vivo* (19). An extra copy of chromosome 6 in *C. parapsilosis* drives cross-tolerance to both tunicamycin and aureobasidin A (41), suggesting that aneuploidy mediates phenotypic changes in this species as well.

However, most LOH that we observed in *C. parapsilosis* started near/at the target locus (Fig. 2), possibly as a result of homology-based repair mechanisms triggered by the Cas9-induced double strand break (DSB), such as gene conversion (short-range LOH) or break-induced repair (long-range LOH) (36, 37). CRISPR-associated LOH has also been observed in *Saccharomyces cerevisiae* (30). Whereas Marton et al (31) suggest that Cas9-associated LOH in *C. albicans* is rare, we note that transformants were selected based on the acquired double prototrophy (Arg+/His+), requiring two <u>H</u>omology <u>D</u>irected <u>R</u>epair (HDR) events with two repair templates, dramatically reducing the likelihood of LOH at the target site. In our experimental design, strains edited at both alleles may be generated by either i) two HDR events, one between each allele and the repair template (RT) (lower risk of LOH), or ii) a single HDR event at one allele, followed by a second recombination between the mutated allele and its homolog (higher risk of LOH).

We also found that targeting the same locus with different guide RNAs can have different effects on LOH. *RHA1* (*CPAR2_805700*) was targeted with two gRNAs (pCP-805700 and pCP-805700-2, Fig. 3). Using pCP-805700 resulted in editing *RHA1* (strain e805700), though the entire gene was not deleted (strain “del”805700) (Fig. 3). Nonetheless, LOH was induced by both attempts (Fig. 2), suggesting that Cas9 did induce a DSB in “del”805700, which was repaired without using the template provided. On the other hand, transformation with pCP-805700-2 resulted in the deletion of *RHA1*, but did not induce LOH (Fig. 3). It is possible that the accuracy of the gRNA, and the efficiency of DSB generation, may influence the level of LOH that occurs.

We found that targeting two different genes on chromosome 8 (*RHA1* and *KIS1*, located approximately 153 and 8 kb upstream of *MET10*, respectively) resulted in LOH that caused cysteine/methionine auxotrophy. Studies from the 1980s using UV treatment to induce mitotic instability in *C. albicans* noted that auxotrophies were common, and cells with a defect in sulfite reduction (auxotrophic for sulfur containing amino acids) were particularly enriched (42-44). In 1998, Whelan and Kwon-Chung (45) described a similar phenomenon in *C. parapsilosis* ATCC 22019 (also known as CLIB214), the parental strain used in this study. We propose that these observations may be explained by UV-induced LOH at *MET10*, which has a stop codon in one allele in *C. parapsilosis* CLIB214.

CRISPR-Cas9-induced LOH in *C. parapsilosis* may muddy the water when interpreting ambiguous phenotypic traits of engineered strains. However, researchers can - and should - have controls in place to avoid associating a mutation with a phenotype. Complementing the mutation introduced is an obvious step to take. Alternatively, LOH could be reduced, by integrating two different nutritional markers in the two alleles (as in (31)). However, this would restrict the use to auxotrophic laboratory strains, and lose many of the benefits of markerless constructs. LOH could also be reduced by using two simple repair templates that differ by a few SNPs, and screening for the presence of both, one at each allele (as in (12)). The 71 heterozygous variants that we identified that could potentially result in deleterious phenotypes as a result of LOH in *C. parapsilosis* CLIB214 constitute a “suspect list” that could be useful in narrowing down a potential culprit for an unexpected phenotype.

## CONCLUSION

We showed that Cas9-induced DSB is often associated with LOH in the opportunistic pathogen *C. parapsilosis*, and that this event can be responsible for phenotypic alterations in the engineered strains. We predicted the existence of at least 12 heterozygous variants (frameshifts and stop codons) in the genome of the strain CLIB214 that are likely to be deleterious following homozygosis. Our findings help define the landscape of unplanned genome changes that the CRISPR-Cas9 technology may induce as collateral effects of gene editing in eukaryotic cells with diploid heterozygous genomes, and stress the importance of confirming a causal link between the introduced mutation and the observed phenotype.

## MATERIALS AND METHODS

### Strains and Media

All *C. parapsilosis* strains (Table S2) were grown in YPD medium (1% yeast extract, 2% peptone, 2% dextrose) or on YPD plates (YPD plus 2% agar) at 30°C. Transformants were selected on YPD agar supplemented with 200 μg/ml nourseothricin (Werner Bioagents, Jena, Germany). Plasmids (Table S3) were propagated in *Escherichia coli* DH5α cells (NEB, United Kingdom) by growing cells in LB media without NaCl (Formedium) supplemented with 100 μg/mL ampicillin (Sigma).

### Phenotypic testing and spot assay

Auxotrophies were identified by growing mutant strains on synthetic complete (SC) dropout media (0.19% yeast nitrogen base without amino acids and ammonium sulfate, 0.5% ammonium sulfate, 2% glucose, 0.075% cysteine/methionine dropout mix, 2% agar). The utilization of different carbon sources was tested on yeast nitrogen base without ammonium sulfate (YNB, 0.19%) supplemented with either 0.5% ammonium sulfate or 10 mM alternative nitrogen sources. For the spot assays, overnight cultures were grown in YPD at 30°C with shaking, washed and diluted to final OD_600 nm_ 0.0625 in 1 ml of PBS. The cultures were serially diluted (1:5) in a 96 well microtiter plate, and then spotted on phenotyping plates with a 48-pin bolt replicator. The plates were grown at 30°C for two days and then photographed.

### Gene editing using pCP-tRNA in *Candida parapsilosis*

The oligonucleotides used to generate and screen the edited mutants are listed in Table S3. The edited strains were constructed by CRISPR-Cas9 editing in *C. parapsilosis* CLIB214 using the pCP-tRNA plasmid ((12), available at Addgene, plasmid # 133812). Suitable guides to induce Cas9 cleavage were computationally designed with EuPaGDT (46), and cloned into the SapI-digested pCP-tRNA plasmid as described in (12). The presence of the guide in the receiving plasmid was confirmed by PCR (M13FWD universal primer + relevant gRNA_BOT oligonucleotide (Table S2)). Repair templates (RTs) for editing (eRTs) were designed to repair the Cas9-induced double strand break (DSB) by <u>H</u>omology <u>D</u>irected <u>R</u>epair (HDR), containing 30 bp homology arms to either side of the cut, 11 bp introducing stop codons in all three reading frames, and a unique 20 bp barcode (tag). For the edited mutant e301940, the RT contains 40 bp homology arms to either side of the cut, two in-frame stop codons (6 bp) and a unique barcode (Table S3). Each eRT was generated by primer extension as described in (12) (Table S3). To delete *CPAR2_805700*, two different 1020-bp long deletion RTs (delRTs) generated by fusion PCR were used, resulting in either: i) the replacement of the entire *ORF* with the barcode, or ii) the partial deletion of a 1248-bp long central region of the *ORF* (+1998/+3245, after amino acid 259) and its replacement with the barcode (Table S3). Three to five μg of purified delRT were used to transform *C. parapsilosis* in combination with the relevant pCP-tRNA plasmid. All oligonucleotides were ordered from Eurofins Genomics. Two independent disrupted strains (A and B) were constructed for each gene target. The names of the edited strains (e.g. e301940) reflect the target gene (e.g. *CPAR2_301940*).

### Transformation of *Candida parapsilosis*

Yeast cells were transformed with 5 μg of the relevant plasmid and either 25 μL of unpurified repair template for editing the gene (eRT) or 5 μg of purified repair template for deleting the gene (delRT), using the lithium acetate method described in (47), with minor modifications (starting OD600nm of YPD culture 0.1 instead of 0.05). Transformants were selected onto YPD agar plates containing 200 μg/mL nourseothricin (Jena Bioscience GmbH, Germany) and screened by colony PCR to confirm the presence of the mutation (Table S2). Representative mutants were sequenced by Sanger sequencing (MWG/Eurofins). For each mutant strain two independent lineages (A and B) were patched onto YPD agar without selection twice to induce the loss of the pCP-tRNA plasmid.

### DNA extraction for whole genome sequencing

The genome DNA of one isolate from each targeted gene disruption/deletion was sequenced. Cells were grown overnight in YPD with shaking, harvested by centrifugation (3,000 rpm, 5 min) and resuspended in 200 μL of Extraction Buffer (2% m/v Triton X 100; 100 mM NaCl; 10 mM Tris pH 7.4; 1 mM EDTA; 1 % m/v SDS) and 200 μL of phenol:chloroform:isoamyl alcohol (25:24:1) (PCIA). Cells were lysed by adding 0.3 g acid-washed glass beads (0.45-0.52 mm, Sigma) and agitating the mixture with a 600 MiniG bead beater (Spex SamplePrep) for 30s (6 times). The mixture was centrifuged (14,000 rpm, 10 minutes), and the aqueous phase transferred in a new tube. The aqueous phase was then extracted by adding 200 μL TE Buffer (pH 8.0) and 200 μL PCIA to the tube and agitating in the bead beater (30s), followed by centrifugation and a second treatment with TE/PCIA, and a third with 200 μL PCIA only.

Nucleic acids were precipitated by adding 1 mL 100% Isopropanol + 80 μL 7.5 M ammonium acetate to the aqueous layer and pelleted by centrifugation. The pellet was washed with 1 mL of 70% ethanol, air-dried, resuspended in 400 μL TE buffer and 1 uL RNAse A (100 mg/mL) and incubated at 37°C for 1 hour. DNA was re-precipitated and washed as above and re-suspended in 100 μL of deionized water. The DNA was then cleaned with the Genomic DNA Clean and Concentrator-10 kit (Zymo Research), according to the manufacturer’s instructions.

### Genome sequencing

Whole-genome sequencing was performed by Beijing Genomics Institute (BGI) on a DNBseq platform generating 150 base paired-end reads. All reads were trimmed using Skewer (v. 0.2.2) (48) to minimal lengths of 30 and average qualities of 35. Trimmed reads were aligned to the *C. parapsilosis* CDC317 reference using BWA mem (v. 0.7.17-r1188) (49). Sorting and duplicate marking was performed using Samtools Sort (v.1.10) and Picard Tools (v. 2.21.6) respectively on output BAM files (50). The Genome Analysis Tool Kit (GATK v. 4.2.0.0) was used to call variants per sample in GVCF format, combining records and joint genotype (51). GATK VariantFiltration was used to filter out variants below a read depth of 15 and minimum genotype quality of 40. Clusters of 5 SNPs in 100 bp windows were removed. Variants flanked by long mono/di-nucleotide repeats were removed using a custom script (https://github.com/CMOTsean/milt_variant_filtration). Heterozygous alleles with a depth ratio of below 0.25 or above 0.75 were also removed (52). To create Fig. 2, sites which were heterozygous in CLIB214 but were homozygous in at least one edited strain were extracted. These sites were plotted as vertical lines along Chromosome 8 using the Matplotlib Python package (53).

Sample coverage and chromosomal copy number were analysed using BEDTools and Delly (54, 55). Annotation of gene variants and prediction of protein coding effects were performed using SIFT. A SIFT4G database for *C. parapsilosis* reference: CDC317 was generated for *Candida parapsilosis* as described in (52, 56).

## Data availability

The data generated in this study have been submitted to the NCBI BioProject database (https://www.ncbi.nlm.nih.gov/bioproject/) under accession number PRJNA866533.

## ACKOWLEDGMENTS

For Open Access, the authors have applied a CC BY public copyright license to any Author Accepted Manuscript version arising from this submission. This study was supported by Science Foundation Ireland (grant numbers 19/FFP/6668 and 18/CRT/6214) and the Irish Research Council (A.R.). We are grateful to Letal I. Salzberg, Eoin Ó Cinnéide, Siobhán A. Turner, and Florent Morio for contributing to the generation of the library of edited strains. We are also thankful to Jeffrey M. Rybak for useful discussion on the manuscript, and to all the members of the Butler and Wolfe’s groups for feedback on the manuscript.

---

**Fig S1.**
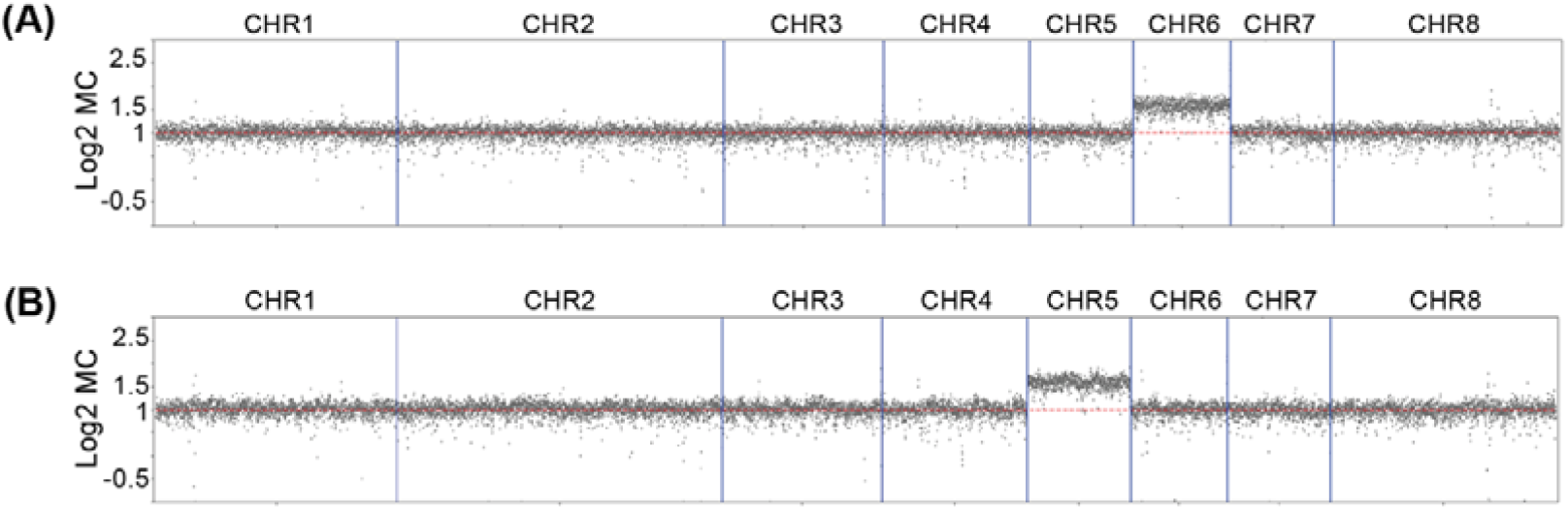
Aneuploidy in CLIB214-derived strains modified using CRISPR-Cas9 technology. Aneuploidy in strains e205070 (A) and e804640 (B), evidenced by Log2 of Mean Coverage (Log2 MC) across the genome. Strains e205070 and e804640 have an extra copy of chromosome 6 and 5, respectively.

**Fig S2.**
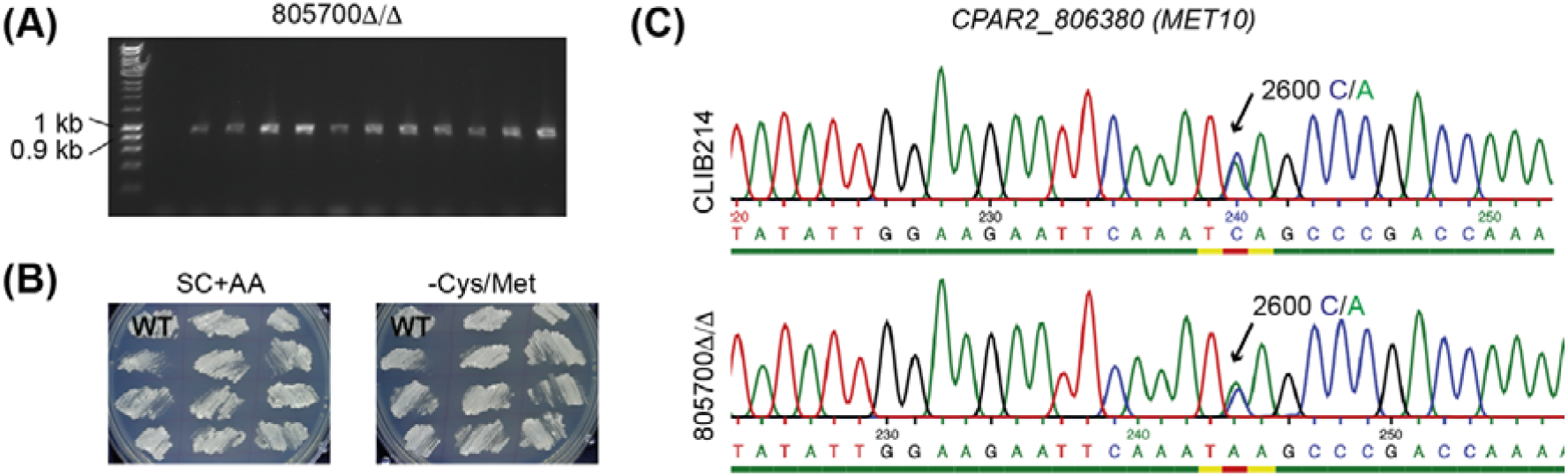
Mutants in which *CPAR2_805700* (*RHA1*) was deleted are cys/met prototrophic. (A) Amplification of the *CPAR2_805700* (*RHA1*) locus confirmed the deletion of the gene (primers s805700-FWD/RhaI-2-dw, Table S3). (B) Growth with (SC+AA) or without (-Cys/Met) cysteine and methionine. CLIB214 (WT) is included as control. All the transformants are prototrophic for cys/met. (C) Amplification and Sanger sequencing of the *CPAR2_806380* (*MET10*) locus in the 805700Δ/Δ isolates confirmed that the heterozygous variant (2600C/A) was maintained. The sequencing traces of CLIB214 and one representative transformant are shown.

**Table S1.**
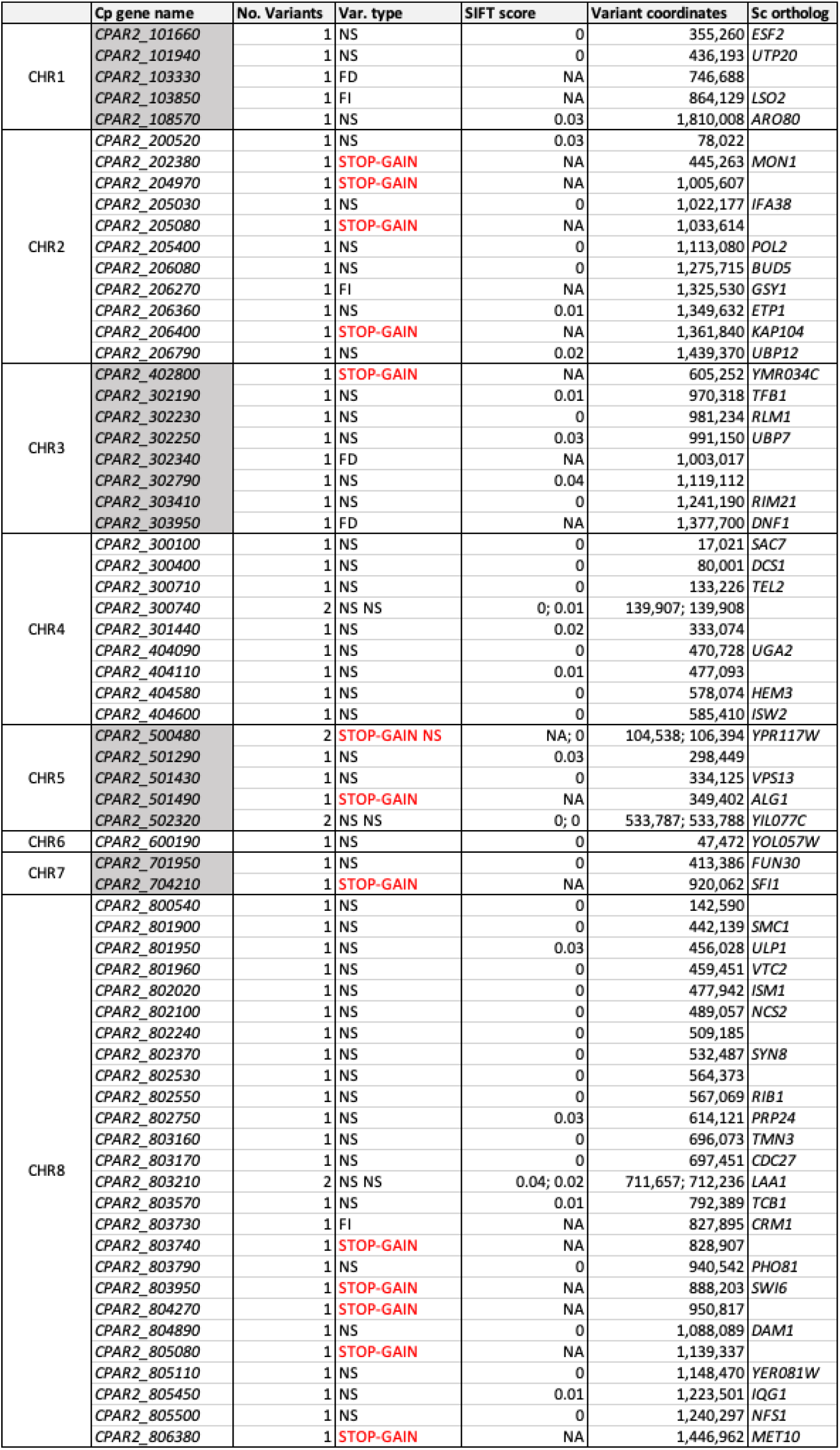
Potentially deleterious heterozygous variants in *C. parapsilosis* CLIB214. FD: frameshift deletion; FI: frameshift insertion; NS: non-synonymous.

**Table S2.** Strains used in this study. The letters A and B represent the two different lineages of a strain.

| Strain | Genotype | Method | Modification of target gene | Reference |
| --- | --- | --- | --- | --- |
| CLIB214 |  |  |  | Type Strain |
| e805700_A | *cpar2_805700::StopTAG/ cpar2_805700::StopTAG* | CRISPR-Cas9 | stop codon | This study |
| e805700_B | *cpar2_805700::StopTAG/ cpar2_805700::StopTAG* | CRISPR-Cas9 | stop codon | This study |
| "del"805700_A | *cpar2_805700/cpar2_805700* | CRISPR-Cas9 | none | This study |
| "del"805700_B | *cpar2_805700/cpar2_805700* | CRISPR-Cas9 | none | This study |
| 805700Δ/Δ_A | *cpar2_805700::TAG/ cpar2_805700::TAG* | CRISPR-Cas9 | deletion | This study |
| 805700Δ/Δ_B | *cpar2_805700::TAG/ cpar2_805700::TAG* | CRISPR-Cas9 | deletion | This study |
| e806320_A | *cpar2_806320::StopTAG/ cpar2_806320::StopTAG* | CRISPR-Cas9 | stop codon | This study |
| e806320_B | *cpar2_806320::StopTAG/ cpar2_806320::StopTAG* | CRISPR-Cas9 | stop codon | This study |
| kis1Δ/Δ | *leu2::FRT/leu2::FRT, his1::FRT/his1::FRT, kis1::LEU2/kis1::HIS1* | HIS1/LEU2 | deletion | (10) |

**Table S3.**
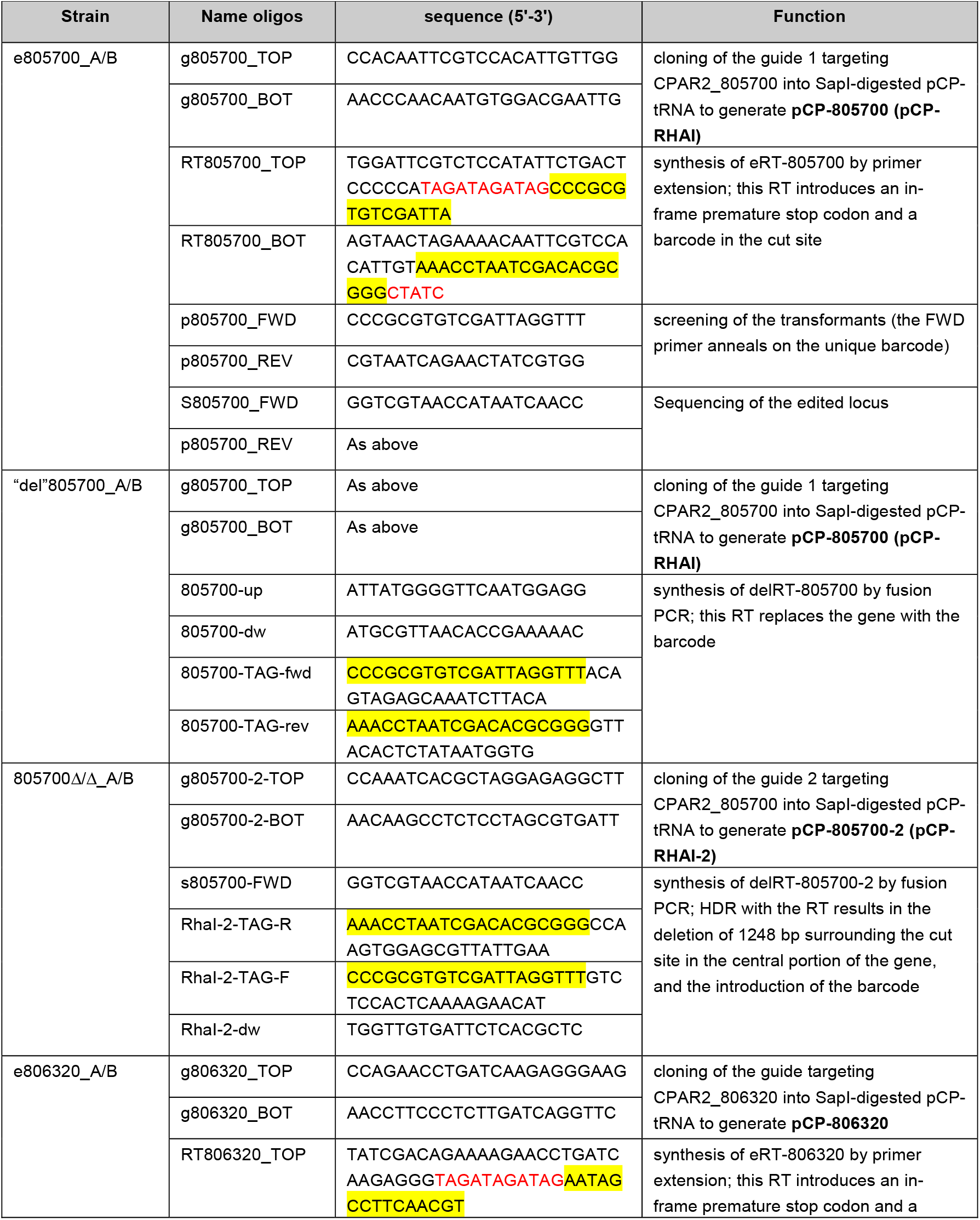
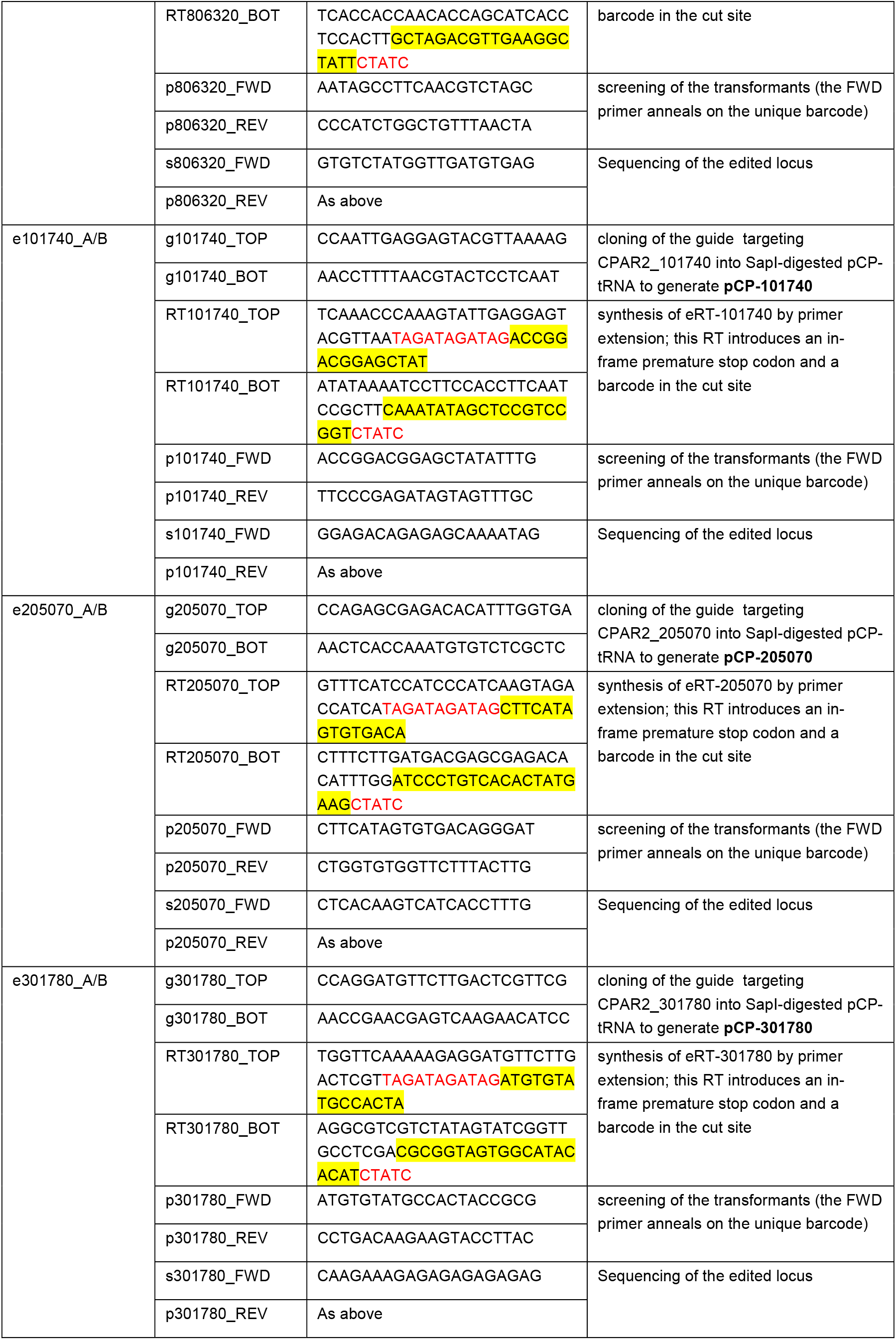
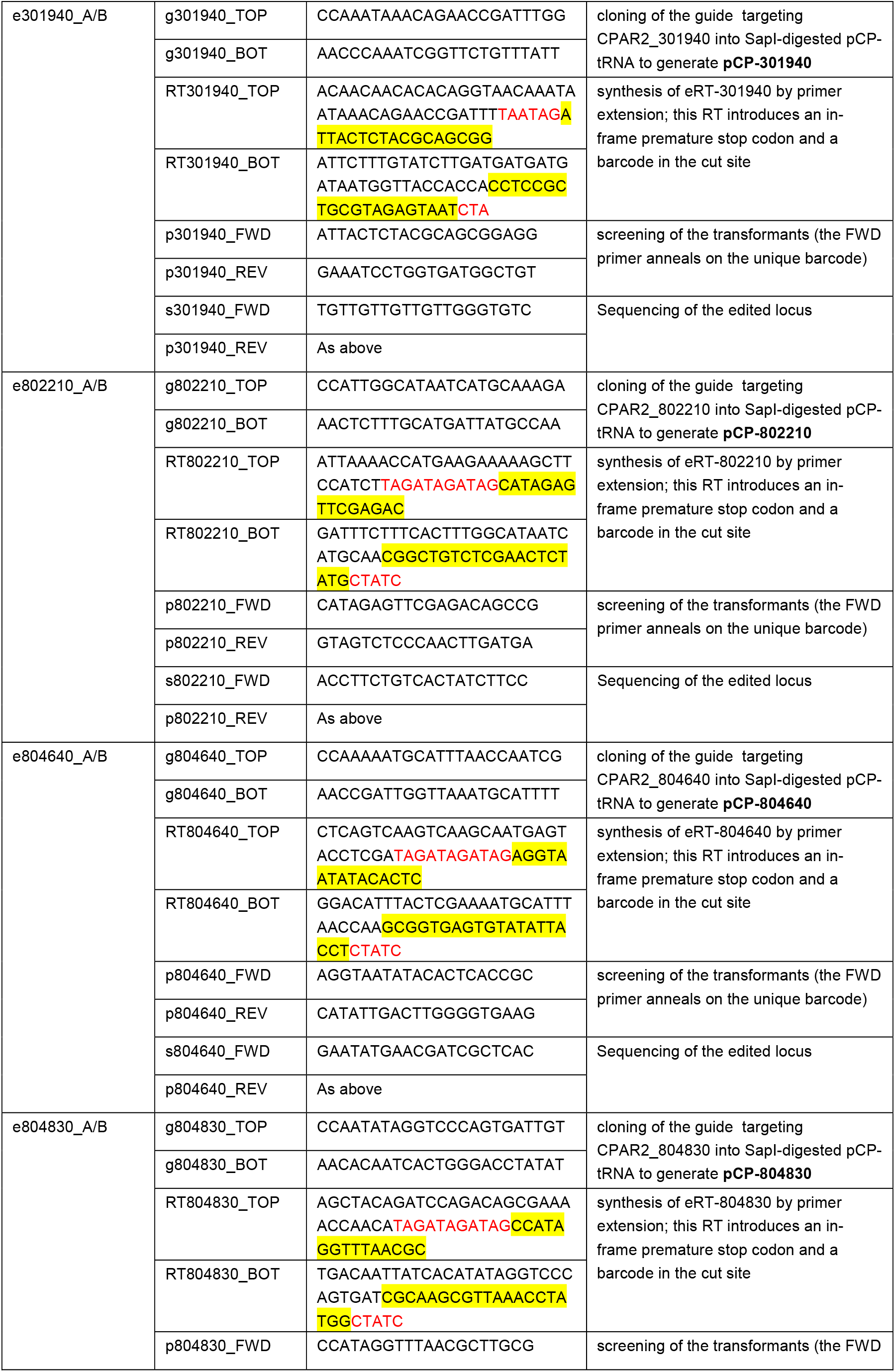
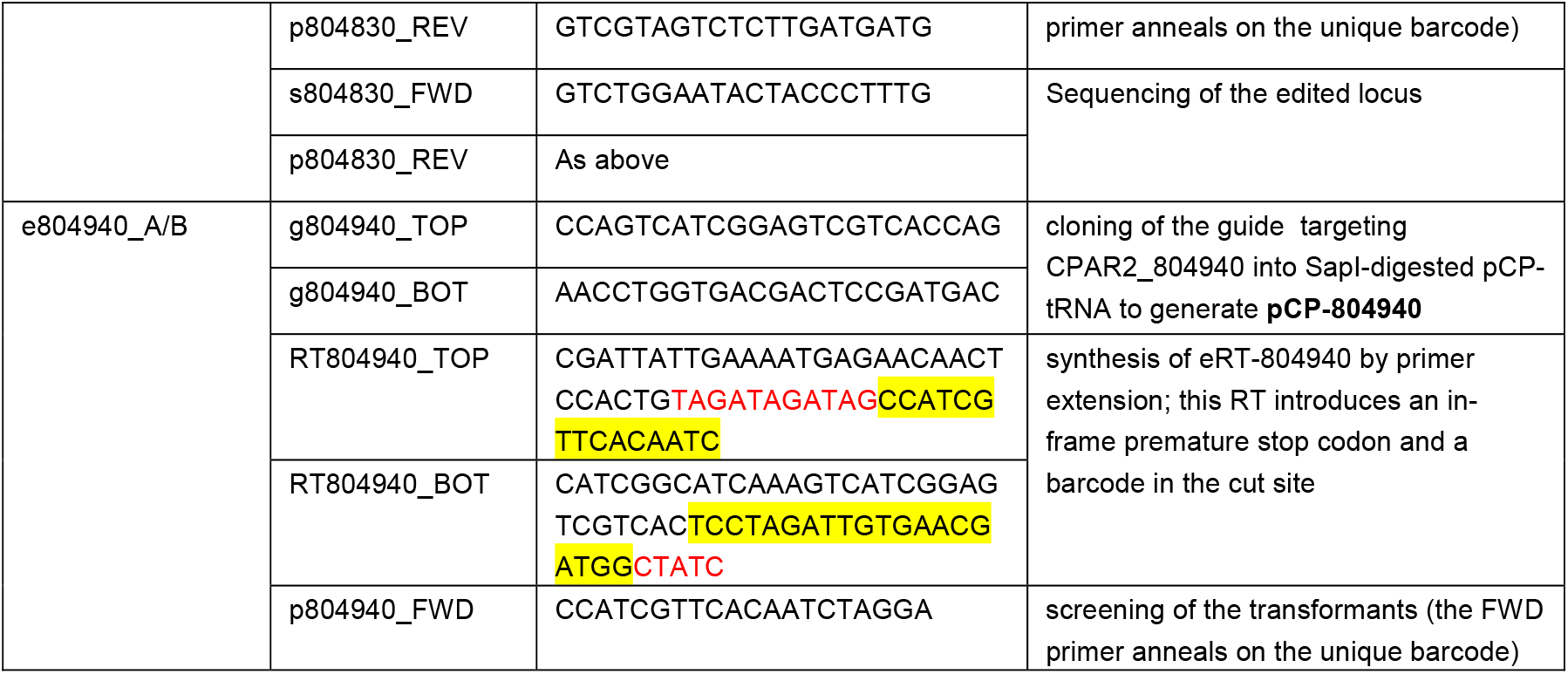
Plasmids and oligonucleotides used in this study. For each strain, the oligonucleotides used for generating the relevant pCP-tRNA plasmid and Repair Templates (RTs) are indicated in the Table, as well as the oligonucleotides used for screening the strains and sequencing the mutated locus. In the sequence of the RT, the barcode and the 11-nt sequence resulting in the introduction of one in-frame stop codon are highlighted in yellow and red, respectively. The plasmids used for introducing the Cas9-mediated cut are indicated in bold. Note that the edited mutant e301940 was generated at an early stage of the construction of the library, and the RT contains two in-frame stop codon (6 bp) and the unique barcode. The primers for sequencing the *CPAR2_806380* (*MET10*) allele in different strains are reported at the end of the table.

